# ContinuumCellAgent: A Framework-Guided Agent for Long-Horizon Scientific Research

**DOI:** 10.64898/2026.06.15.732409

**Authors:** Hao Li, Yifei Lu, Kaiwen Fang, Zixi Xu, Fuhai Li

## Abstract

AI-scientist systems are beginning to automate parts of scientific research. We present ContinuumCellAgent, an autonomous agent that executes literature review, hypothesis formation, computational experimentation, manuscript drafting, and adversarial peer review as a single unattended run. Existing AI scientist systems remain difficult to diagnose because they lack modularity, systematic prompt grounding, and observability into long-running behavior. ContinuumCellAgent addresses these gaps with a modular supernode architecture for stage-wise backend swapping, protocols grounded in curated research-method checklists that also define reviewer rubrics, and a diagnostics layer that records file-based artifacts, message traces, and state transitions. We evaluate the system on open-domain QA benchmarks and biomedical/longevity case studies, showing that it can produce checkable research artifacts while exposing pipeline dynamics for rigorous AI co-scientist research.

## 1 Introduction

The vision of an AI system that autonomously conducts scientific research— formulating hypotheses, designing experiments, executing code, and writing papers—has motivated a growing body of work [3, 5, 13, 19]. Recent advances in large language models have made this vision increasingly tangible: agents can now search literature, write and debug code, and produce coherent manuscripts. Yet the path from demonstration to rigorous, reproducible autonomous science remains fraught with under-explored challenges.

Existing systems typically automate only one or two stages of the research lifecycle. *ResearchAgent* [3] generates ideas via knowledge-graph traversal and LLM refinement but does not execute experiments or produce manuscripts. *MLAgentBench* [17] benchmarks coding agents on machine-learning tasks but lacks literature review, hypothesis generation, and paper-writing stages. *The AI Scientist* [19] covers the full cycle— ideation, coding, writing, and self-review—but operates exclusively on ML-template experiments with pre-written scaffolding, uses a monolithic architecture that cannot swap agent backends, and provides no diagnostics into *where* or *why* the pipeline fails.

We identify three gaps that limit progress toward robust autonomous scientific discovery:

### G1. Modular agent composition

No existing system lets researchers plug different agent backends into different stages of the same pipeline. This makes it impossible to perform controlled ablations—e.g., “Does a planner-executor outperform a simple ReAct agent at modeling when ideation and paper-writing are held constant?”—or to leverage provider-specific strengths (one model/agent/prompt may excel at retrieval, another at code generation). The lack of modularity also impedes extensibility: adding a new agent architecture requires rewriting the entire system rather than swapping a single component. As the landscape of LLM agents evolves rapidly, this rigidity becomes a significant barrier to systematic research.

### G2. Systematic prompt grounding

Agent prompts are typically written ad hoc, reflecting the intuitions of individual developers rather than established methodological principles. Yet decades of scholarship have codified best practices for scientific reasoning (Pearl’s causal hierarchy [21], Popper’s falsifiability [22], Chamberlin’s multiple working hypotheses [7]), statistical rigor (ASA Statement on *p*-values [26], Gelman’s forking-paths analysis [8]), and scientific writing (Schimel’s OCAR structure [24], Toulmin’s argument model [25]).

We use these frameworks to make agent behavior more disciplined and—crucially—to provide the adversarial reviewer with concrete, auditable criteria. When instructions and evaluation rubrics derive from the same source, we eliminate the “misalignment tax” where agents optimize for implicit criteria different from those used to judge them. The prompts can be studied through benchmark results, but they are also part of the explanation for why an agent reported a claim, rejected a result, or revised a manuscript. Prior systems such as FARS [1] and The AI Scientist [19] discuss paper acceptance or final quality as evaluation signals; we argue that the process of making the science, including the quality of intermediate reasoning, should also be inspectable.

### G3. State-level observability and multi-phase evaluation

Prior work reports final output quality (e.g., paper acceptance rates, benchmark scores) but not *where in the pipeline* the agent struggles or *how* revision loops affect quality over time. As autonomous research systems become longer-running and more complex, final-outcome metrics alone are insufficient. We need fine-grained observability: Which stage consumes the most time? Where do failures cluster? Do additional revision iterations improve quality, or do they risk context-window exhaustion? Furthermore, evaluation itself is multifaceted: automated benchmarks with known answers test correctness; AI reviewers provide scalable quality estimation; and expert case studies validate real-world utility. No prior system integrates all three evaluation modalities.

ContinuumCellagent addresses all three gaps. Its contributions are:

### C1. Modular Supernode Architecture (Section 3)

We organize the research lifecycle as a LangGraph state machine with three *supernodes*—Ideation, Modeling, and Paper—each defined by a protocol, a tool set, message memory, file artifacts, and an iteration budget. Each supernode is followed by an adversarial *review gate* that can accept (advancing to the next stage) or reject and route back to *any* upstream stage. Crucially, the agent backend powering each supernode is a configuration choice: command-line flags such as `--modeler` `react` and `--modeler planner-executor` swap the Modeling backend, enabling controlled experiments. This modularity also facilitates *agent-level hyperparameter optimization*—an under-explored research direction where the “hyperparameters” are architectural choices like agent type, iteration budget, and tool set.

### C2. Framework-Grounded Protocols (Section 3.2)

We compiled operationalizable methodological frameworks into a reference document and encoded the relevant subset as stage-specific checklists embedded directly in agent protocols. For Ideation: Chamberlin’s multiple hypotheses, Popper’s falsifiability, Pearl’s causal ladder, Heilmeier’s catechism. For Modeling: Tukey’s EDA protocol, Gelman’s forking-paths documentation, ASA’s statistical rigor. For Paper: Schimel’s OCAR narrative, Toulmin’s argument structure, Feynman’s integrity principle. The same checklists serve as evaluation rubrics for the adversarial reviewer, ensuring alignment between agent instructions and assessment criteria. Every stage ends with a Gawande-style DO-CONFIRM exit checklist that catches systematic omissions before yielding control.

### C3. State Dynamics Diagnostics & Multi-Phase Benchmarking (Section 3.3, Section 4)

A built-in diagnostics layer emits structured timeline events (`stage_start`, `stage_end`) to a JSONL file, recording per-stage wall-clock time, iteration counts, and outcomes. Analysis scripts produce stage-level profiling, Gantt-style timelines, and failure-mode taxonomies. We evaluate ContinuumCellAgent with biomedical/longevity case studies and open-domain QA benchmarks, and use the traces to study stage timing, rollback depth, and failure modes.

The remainder of this paper is organized as follows. Section 2 surveys related work and positions ContinuumCellAgent within the landscape of autonomous scientific agents. Section 3 details the supernode architecture, framework-grounded protocols, traceable execution, session isolation, and LLM fallback chain. Section 4 reports biomedical/longevity case studies and open-domain QA results. Section 5 discusses what the current evidence does and does not establish, and Section 6 concludes.

## 2 Related Work

Table 1 positions ContinuumCellAgent against the most relevant prior systems along six capability axes.

**Table 1:**
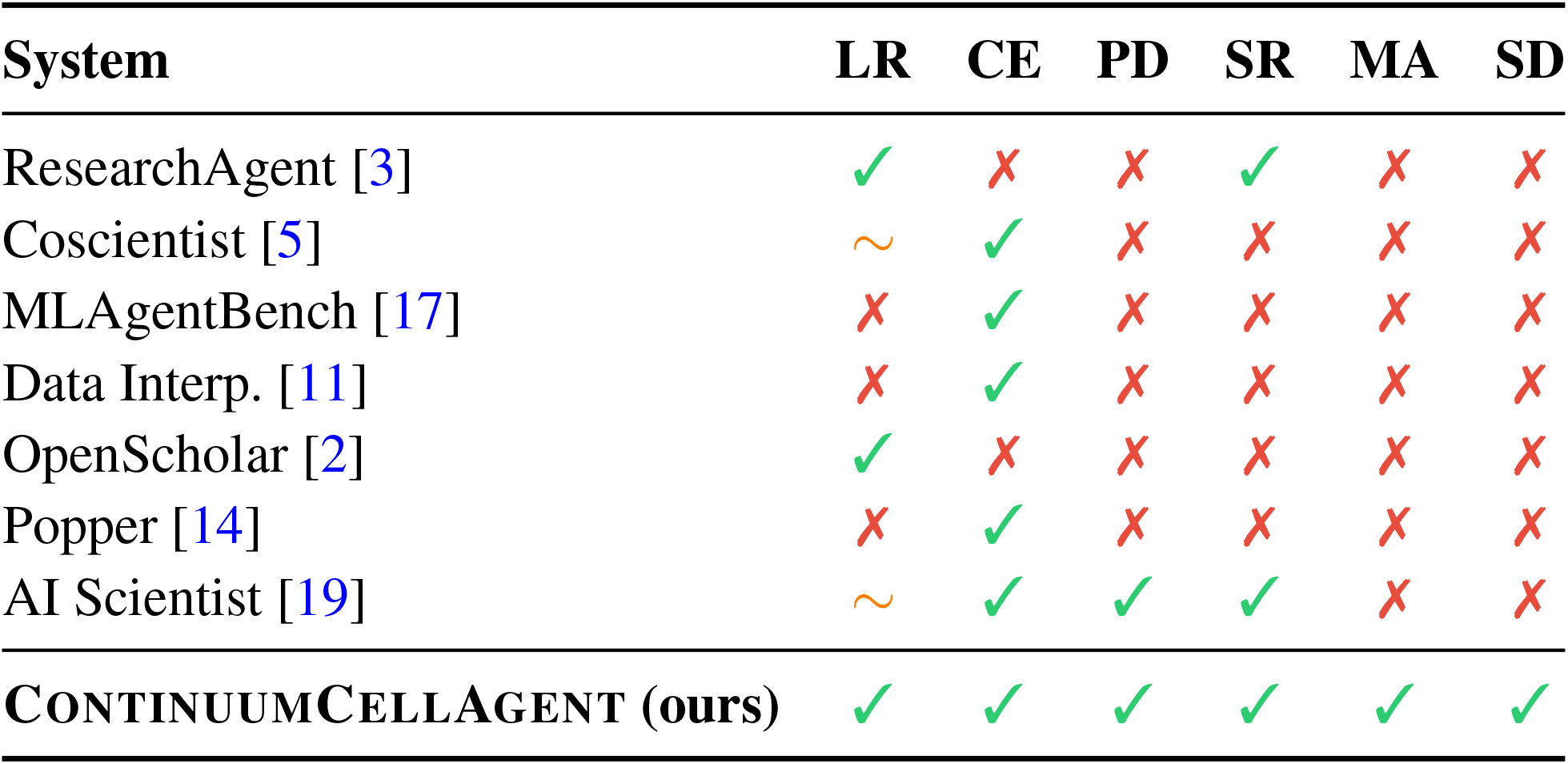
Comparison of autonomous scientific-discovery systems. 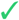 = supported, 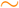 = partial, 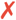 = absent. Columns: Lit. Review (LR), Code Exec. (CE), Paper Draft (PD), Self-Review (SR), Modular Agents (MA), State Diag. (SD).

### Ideation or search-only systems

ResearchAgent [3] uses an academic knowledge graph to propose research ideas, iteratively refined by LLM reviewers. OpenScholar [2] retrieves and synthesizes literature with RAG over 45M papers. Neither executes experiments.

### Code-execution agents

MLAgentBench [16] and Data Interpreter [11] excel at coding tasks but lack literature review and manuscript production. Popper [14] validates hypotheses via falsification experiments but does not generate hypotheses or write papers.

### Domain-specific agents

ChemCrow/Coscientist [5] integrates chemistry tools (RDKit, robotic labs) but is single-domain and produces no manuscript. In biomedicine, OmniCellAgent [12] and Biomni [15] integrate domain databases and specialized analysis tools, but their validation remains field-specific and their designs are less suited to open-ended, end-to-end scientific studies. Biomedical case studies are a common application for such systems: Robin automates a path from hypothesis generation to analysis, but its validation is case-study based and its components are not designed for fine-grained self-evolution or modular backend swaps. Kosmos [20] extends the scope to a broader range of open- and closed-ended problems. BioMedAgent [6] adds a self-evolving mechanism, but focuses on expert-curated biomedical analysis questions rather than an end-to-end research workflow that produces a manuscript.

### Full-cycle systems

The AI Scientist [19] is the closest predecessor: it generates ideas, writes code, runs experiments, drafts papers, and self-reviews. However, it (i) operates only on ML-template experiments with pre-written scaffolding, (ii) uses a monolithic architecture with no mechanism to swap agent backends per stage, and (iii) provides no state-level diagnostics. ContinuumCellAgent generalizes the full-cycle approach to arbitrary scientific domains (demonstrated on biomedical research), introduces modular agent composition, and adds a diagnostics layer for studying agent dynamics.

## 3 Method

ContinuumCellAgent is designed as practical research software rather than a single monolithic prompt. Its main structure is a **LangGraph state machine** with three major tasks—Ideation, Modeling, and Paper—implemented as *supernodes*. Each supernode owns a stage-specific protocol, tool set, working memory, and file-based artifacts, and each is followed by an adversarial review gate that can accept the result or route execution back to any previous stage (Figure 1). This design makes the pipeline useful both as an autonomous research assistant and as an experimental harness: the same task can be rerun with different agents, models, budgets, or tools while preserving the protocol and trace format.

**Figure 1:**
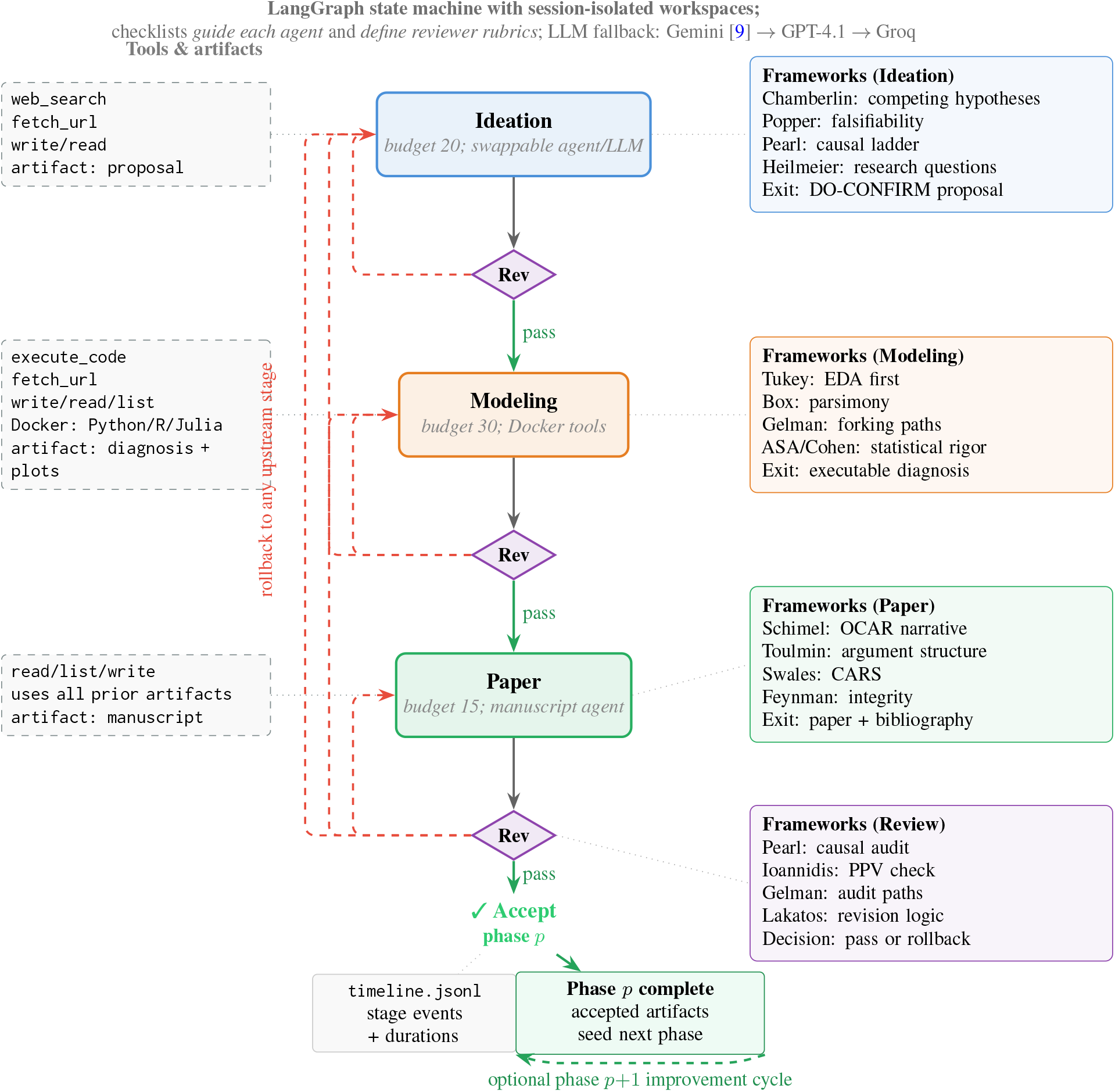
Top-down annotated flowchart of ContinuumCellAgent. The center spine shows the three supernodes (Ideation, Modeling, Paper)—each with a swappable agent backend and LLM—interleaved with adversarial review gates that either pass forward (green) or roll back to any upstream stage via the dashed red lanes on the far left. A final accept closes one research phase; the accepted artifacts can seed a later phase, shown by the dashed green loop under the phase-complete block, enabling multi-phase improvement rather than treating acceptance as the end of all work. The left column lists the tool set bound to each supernode (only Modeling uses the Docker sandbox); the right column lists the methodological frameworks that ground each stage’s protocol and simultaneously serve as the reviewer’s rubric.

**Figure 2:**
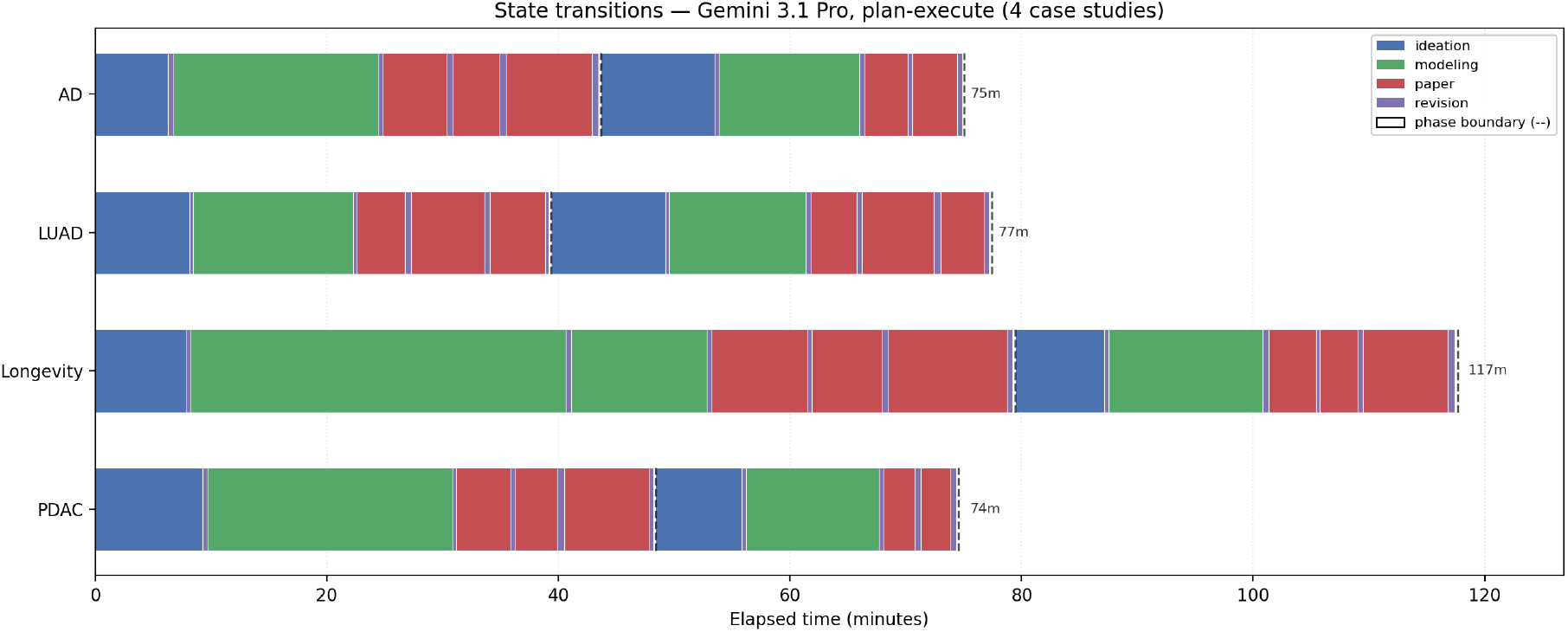
State-transition summary across four provider-LLM Plan-Execute case studies (AD, LUAD, Longevity, and PDAC). Each horizontal bar shows elapsed time through Ideation, Modeling, Paper, and review gates, with dashed markers denoting phase boundaries. The aggregate view makes execution policy inspectable at a level that final papers and scalar scores hide: forward progress, reviewer-triggered rollbacks, and repeated stages across cases. We use this diagnostic as descriptive evidence about agent dynamics, not as a standalone quality metric.

### 3.1 Supernodes and Review Gates

Each supernode is defined by a *protocol*, not by a particular LLM provider. The protocol specifies the stage objective, allowed tools, memory interface, required output files, iteration budget, and exit checklist. Ideation searches literature and writes a proposal with competing hypotheses; Modeling writes and executes Python/R/Julia code in a Docker sandbox and reports statistical results; Paper reads all accumulated artifacts and drafts a manuscript with citations. Full per-stage tool specifications, budgets, session isolation, and the LLM fallback chain are in Appendix A.

The stages use both **file-based state** and **message-based memory**. File tools read and write durable artifacts such as `research_proposal.md`, source code, plots, `diagnosis_report.md`, manuscripts, and bibliographies. The message state carries the current task, prior reviewer decisions, summaries, and routing context across the LangGraph. This separation is important in practice: files make scientific outputs inspectable and rerunnable, while memory lets the agent coordinate long-horizon decisions without re-parsing the whole workspace at every step.

The **review gates** are lightweight, tool-free LLM calls that fire after every supernode. Each gate scores the stage’s output on stage-appropriate criteria—gap validity for ideation, statistical rigor for modeling, novelty and soundness for the paper—and returns either ACCEPT or a routing directive to an upstream stage. Because the gates use the same framework-grounded rubrics embedded in the supernode protocols (Section 3.2), agent instructions and evaluation criteria stay aligned.

### Swappable agents and adapters

Because a supernode exposes a stable protocol—inputs, tools, required artifacts, review rubric, and trace schema—the implementation can swap the backend agent without changing the rest of the pipeline. A command-line flag (e.g., `--modeler react` or `--modeler planner-executor`) can replace the Modeling backend while holding ideation, paper writing, tools, and review budget fixed. The same adapter pattern generalizes to other stages and enables heterogeneous configurations, prompt-adjustment variants, and agent-level hyperparameter optimization over choices such as backend, prompt variant, iteration budget, and tool set.

### Prompt-based evolution

Plan-Evolve is a prompt-adjustment variant of Plan-Execute. It starts from the same initial stage prompt, runs the agent with the available tools, and then lets the agent revise the prompt text as the run proceeds. Subsequent attempts therefore use adjusted prompts rather than a separate training procedure. The adapter interface remains the same for comparing ReAct, Plan-Execute, and Plan-Evolve.

### 3.2 Framework Grounding as Protocol Design

The protocol for each supernode is grounded in methodological checklists rather than ad hoc instructions. We compiled operationalizable frameworks into a reference document (`FRAMEWORKS_REFERENCE.md`, 1705 lines) and encoded the relevant subset as stage-specific checklists embedded directly in agent protocols.

The right column of Figure 1 summarizes the mapping from frameworks to pipeline stages and shows that the same checklists drive both agent execution and reviewer evaluation.

#### Dual-use design

A checklist serves two roles simultaneously: (1) it *guides* the agent during execution (“have I classified my hypothesis on Pearl’s causal ladder?”), and (2) it provides the *reviewer* with concrete criteria to audit (“did the paper overclaim causation from Rung-1 evidence?”). Because the review gate fires after *every* supernode, each stage’s output is validated before downstream stages consume it. This eliminates the common failure mode where agent and reviewer operate under different implicit standards, and catches errors early (e.g., a flawed hypothesis is rejected before expensive modeling begins).

#### DO-CONFIRM exit protocol

Inspired by Gawande’s *Checklist Manifesto*, every supernode works freely during its iteration budget, then runs a mandatory exit checklist before yielding control. This catches systematic omissions (e.g., missing effect sizes, unreported negative results) without constraining the agent’s exploratory freedom during main execution.

### 3.3 Traceable Execution and Agent Dynamics

For long-horizon agent evaluation, the intermediate trajectory is as important as the final artifact. Continuum-Cellagent therefore logs both durable scientific artifacts and structured execution traces. Every session uses an isolated workspace with a concrete folder layout so that sources, generated code, figures, diagnosis reports, reviewer messages, and manuscripts can be inspected after the run. In parallel, every node execution emits structured events to a `timeline.jsonl` file:

- `stage_start`: stage name, wall-clock timestamp, iteration index.
- `stage_end`: duration in seconds, outcome (`completed` or `failed`), iteration index.

A companion analysis script (`analyse_timeline.py`) produces: (i) a per-stage breakdown table with wall-clock time and percentage, (ii) a Gantt-style timeline plot, and (iii) a structured JSON summary.

This traceability is what makes the experiments in this paper possible. The saved sources and executed code make case-study claims checkable, while the state trace supports analyses of time-to-transition, rollback depth, score-versus-cost tradeoffs, and where the system fails or improves under revision. We view these analyses as an opening for future agent-dynamics work rather than a solved evaluation protocol.

### Stage profiling

The timing trace is an auditing device rather than a deterministic claim about how long each stage must take. It lets us inspect where a particular controlled run spent time, provided the run configuration is kept visible: per-stage step budgets, maximum revision count, selected run phases, backend latency, tool runtime, and data availability all affect the observed allocation. In the reported case studies, modeling is the largest wall-clock component, but we treat that as a descriptive property of these runs under fixed controls, not as an intrinsic law of the workflow.

### Failure-mode taxonomy

We observe three primary failure modes: (1) *context-window exhaustion* after extended revision loops (the accumulated conversation exceeds provider limits), (2) *sandbox execution errors* (missing packages, data download failures), and (3) *reviewer hallucination* (the reviewer requests changes already present in the manuscript). Mode (1) is the most common and suggests a need for context summarization between iterations.

### Revision productivity

Not all reviewer-triggered revisions are productive. Our diagnostics let us inspect how reviewer scores and failure risk change as the run approaches its configured revision cap. In the current runs, early revision cycles tend to improve the manuscript, while later cycles show diminishing returns and greater context-window risk; this motivates treating max revisions and enabled run phases as experimental controls rather than assuming an open-ended revision process is always better.

## 4 Experimental Setup and Results

### 4.1 Evaluation Questions and Setup

We evaluate three questions that match what the current system can support: (Q1) Can a protocol-grounded agent complete useful end-to-end scientific workflows with saved sources, executable code, and reported artifacts? (Q2) Can the same framework also support open-source question-answering benchmarks, showing that the design is not only a biomedical demo? and (Q3) Does the traceable implementation make the agent more inspectable—revealing stage transitions, revision depth, time cost, and failure modes—even when those diagnostics raise new questions rather than closing them? To answer them, we combine biomedical case studies with open-domain QA benchmarks and trajectory logs.

### Models

For the open-source QA tests, the no-tool baseline and all ReAct, Plan-Execute, and Plan-Evolve agents use the same locally hosted Qwen3.5-27B model [23] served with `sglang`. For the Tier-1 case studies, ContinuumCel-lAgent uses provider LLMs through a common adapter layer: the case-study traces use Plan-Execute trajectories from Gemini 3.1 Pro [9] and GPT-family configurations. Serving and sandbox-package details are given in Appendix A.

### Tier 1: End-to-end biomedical discovery (reported in this paper)

1. DAC-research: “Identify key dysfunctional genes and disrupted pathways in pancreatic ductal adenocarcinoma (PDAC). Perform differential expression and pathway enrichment analysis on public RNA-seq data, report top dysregulated genes, enriched GO/KEGG terms, and a publication-ready volcano plot.”
2. AD-research: “Investigate dysregulated genes and molecular pathways underlying Alzheimer Disease. Analyze public transcriptomic datasets, perform differential expression and gene-set enrichment, and report candidate therapeutic targets with supporting statistical evidence.”
3. LUAD-research: “Characterize dysfunctional genes and pathway alterations in lung adenocarcinoma (LUAD). Conduct differential expression analysis on TCGA or GEO data, identify driver pathways via enrichment analysis, and benchmark against known LUAD driver genes from the literature.”
4. Longevity-research: “Find the molecular targets and signaling pathways and other factors of longevity. Survey the literature and public transcriptomic / multi-omics datasets across long-lived models (e.g. centenarians, naked mole rat, caloric-restricted cohorts), identify high-confidence longevity-associated genes and pathways (IGF1/mTOR, AMPK, sirtuins, autophagy, senescence, NAD+ metabolism, etc.), perform differential expression or set enrichment where data permits, and report ranked candidate targets, supporting statistical evidence, and a publication-ready visualisation.”

### Tier 2: Open-domain retrieval and multi-hop QA

To test retrieval reliability, multi-hop reasoning, and answer faithfulness beyond the biomedical case studies, we run the same agent on TriviaQA, HotpotQA, and 2WikiMultihopQA from LongBench v1 [4], with task counts chosen for available compute and evaluated across the available agent strategies (ReAct, Plan-Execute, Plan-Evolve) with exact coverage accounting (Section 4.3):

1. **TriviaQA, HotpotQA, 2WikiMultihopQA** (LongBench v1 English retrieval-QA subsets; 200 questions per subset for ReAct and Plan-Execute): single- and multi-hop factoid QA with published 30B-class baselines.

Broader agentic-environment benchmarks that require live browser or desktop actuation, including GAIA, WebArena, VisualWebArena, OSWorld, and AgentBench, fall outside our current text-and-retrieval tool set and are left to future work (Appendix B).

### Tier-2 sampling protocol

Coverage is reported per benchmark and agent. ReAct and Plan-Execute complete the 200-question evaluation sample for each selected LongBench subset; Plan-Evolve is evaluated on its deterministic 120-question run per subset with prompt adjustment enabled. The selected tasks are recorded so that any strategy comparison can be interpreted with its exact coverage.

Unless noted otherwise, Tier-1 biomedical runs reported in the main text use Gemini 3.1 Pro as the primary LLM, max 5 revision iterations, and the Docker scientific stack described in Appendix A.

### 4.2 Main Results

#### End-to-end capability and checkability

AD and LUAD complete the full research pipeline across five revision iterations. PDAC completes four full cycles and fails on the fifth modeling invocation because accumulated context exceeds provider limits. Each case saves sources, executable Python/R code, plots and metrics, reviewer feedback, a diagnosis report, and a manuscript with bibliography. We therefore treat completion as an auditable artifact bundle, not as evidence that the scientific conclusion is correct.

### Failure modes and traces

The same artifacts expose scientific failures. In the AD run, the manuscript claims real-world cohorts although the executed pipeline used generated synthetic data; in PDAC, the agent acquired GEO data but left pathway-enrichment errors and stromal-contamination concerns unresolved. Stage logs show modeling as the largest time component in these runs and review as a rollback trigger, but these patterns are descriptive under fixed controls (revision caps, enabled phases, stage budgets, backend latency, and tool runtime), not deterministic properties of the tasks.

### 4.3 Tier 2: Open-Domain QA Results

We evaluate locally hosted Qwen3.5-27B with no tools, with three agent strategies, and against comparable open-source anchors. Pure-model LongBench anchors come from the LongBench work and released source code [4]; Search-R1/R1-base anchors are reported in Search-R1 [18], with the R1 baseline tied to Guo et al. [10]. The selected LongBench rows are provided-context F1 anchors, whereas our agents receive only the question and retrieve evidence by open-web search. Search-R1 is a question-only EM anchor, but in a controlled Wikipedia-2018/E5 retrieval environment. We report F1_NP_ for LongBench-style single-span comparison and EM for question-only QA anchors.

#### TriviaQA, HotpotQA, and 2WikiMultihopQA

Figure 3 shows that all agent variants improve over no-tool Qwen on recall and stricter answer-span scoring; even on TriviaQA, where baseline recall is already high, recall increases slightly. As zero-shot agent loops without benchmark-specific RL training, the best agent variants substantially improve over the raw Qwen baseline and approach the separate trained-search track despite not receiving LongBench supporting passages. The raw-model F1 values are conservative because the scorer does not strip Qwen’s emitted thinking process before final-answer scoring; a Qwen-specific parser may be needed for calibrated no-tool F1. Against provided-context LongBench anchors, the best agent variants exceed both open-source rows on 2WikiMultihopQA and are mixed on the other subsets. Plan-Evolve tests whether prompt adjustment during the run helps; in these results it is not a clear improvement over ReAct.

**Figure 3:**
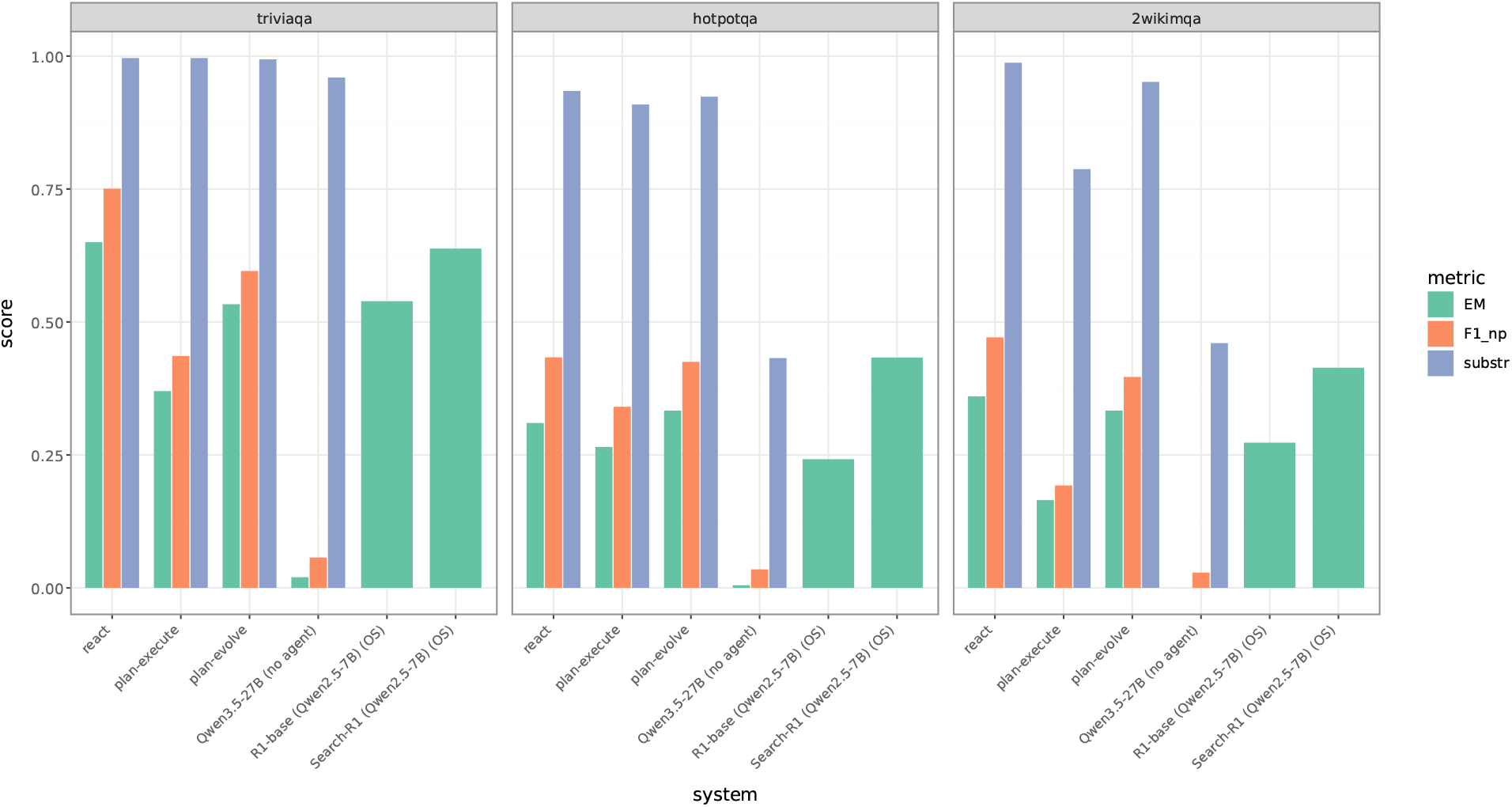
Tier-2 results on TriviaQA, HotpotQA, and 2WikiMultihopQA: each agent strategy (ReAct, Plan-Execute, Plan-Evolve) vs. the no-tool Qwen3.5-27B baseline and published anchors, faceted by benchmark and grouped by metric.

**Figure 4:**
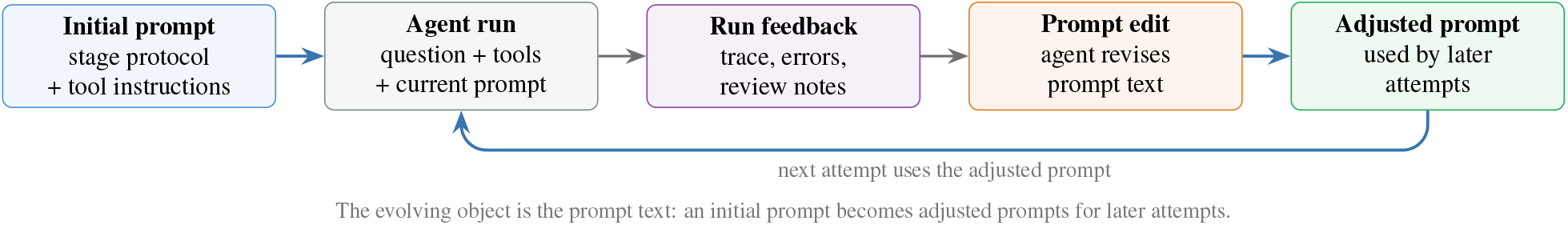
Appendix view of Plan-Evolve prompt adjustment. The agent starts from an initial stage prompt, observes run feedback, edits the prompt, and uses the adjusted prompt in later attempts.

### 4.4 Controlled Backend Comparison

The adapter design supports controlled backend comparisons by fixing the prompt, workspace, tools, reviewer rubric, and trace schema while the agent strategy changes. Tier 2 uses this for ReAct, Plan-Execute, and Plan-Evolve; a paired biomedical backend ablation remains future work.

## 5 Discussion

The experiments support a narrow claim: protocol-grounded tool loops can produce inspectable biomedical artifacts and improve a Qwen3.5-27B baseline on TriviaQA, HotpotQA, and 2WikiMultihopQA (Figure 3). They do not establish autonomous scientific validity. The AD and PDAC cases show why: a run can finish while using synthetic data or leaving core analysis errors unresolved. The main value is therefore auditability—linking final outputs to sources, code, state transitions, and reviewer decisions. Timing and rollback traces are instrumentation under a fixed run configuration, not general metrics. Backend swapping and Plan-Evolve are useful experimental variables only with matched prompts, tools, and revision budgets.

## 6 Conclusion

We presented ContinuumCellAgent, a traceable testbed for protocol-grounded scientific agents. In the current experiments, it saves code, sources, intermediate states, and reviewer decisions; it also improves a no-tool Qwen3.5-27B baseline on TriviaQA, HotpotQA, and 2WikiMultihopQA (Figure 3). The caution is equally important: workflow completion is auditable, but it is not scientific validity. Reliable autonomous science will require stronger provenance checks, paired backend ablations, and domain-expert review of the biological claims.

## Limitations

The current system remains limited by coupled failure modes common to long-horizon scientific agents: context-window exhaustion, unstable memory, brittle biomedical data acquisition, mismatches between cited and processed datasets, shallow recovery from execution errors, and occasional substitution of synthetic or prototype data when real data access fails. Benchmark results are likewise conditional on task structure, coverage, metric choice, and judging protocol; the Plan-Evolve rows are also conditional on the prompt-adjustment trajectory used in the run, and this version reports TriviaQA, HotpotQA, and 2WikiMultihopQA rather than the broader LongBench suite. Finally, automated reviewers are useful for scalable triage but cannot replace domain-expert review for scientific validity, especially when data provenance or biological interpretation is at stake.

## A Implementation Details

### Model serving and providers

Tier-2 QA runs serve Qwen3.5-27B locally through `sglang`; the same served model is used for the no-tool baseline and all agent loops. Tier-1 case-study runs call provider LLMs through the same adapter layer: the reported Plan-Execute trajectories include Gemini and GPT-family configurations.

### Per-stage tool specifications

Each supernode binds a small, stage-specific tool set. Ideation (20-step budget) can search, fetch, read, and write, and produces `references/research_proposal.md`. Modeling (30-step budget) adds `execute_code` and `list_files`, producing `code/results/diagnosis_report.md`. Paper (15-step budget) uses read/list/write tools over prior artifacts and produces `paper/manuscript.md`.

### Session isolation

Every run creates an ephemeral workspace under `instances/<session>/`. All file I/O tools resolve paths relative to this workspace and reject path-traversal attempts. Code executes in a Docker container (`continuum-sandbox`) with NVIDIA CUDA 12.4, pre-installed Python 3.10 packages (PyTorch, scikit-learn, Scanpy, NumPy, SciPy, pandas, statsmodels, Matplotlib, and Seaborn), R 4.3 packages (DESeq2, Seurat, and clusterProfiler), and Julia 1.11. Only the session folder is mounted; the host is never exposed.

### LLM fallback chain

ContinuumCellAgent constructs a LangChain `.with_fallbacks()` chain from available API keys: Gemini *→* OpenAI GPT-4.1 *→* Groq. If the primary provider returns a 429 or 5xx, the request transparently retries on the next provider, supporting unattended operation without making availability deterministic.

## B Cross-Domain Benchmark Suite (Tier 2)

To evaluate behavior beyond biomedical case studies, Tier 2 focuses on retrieval-QA tasks that stress retrieval fidelity, multi-hop reasoning, answer extraction, and cross-domain transfer.

### Benchmarks evaluated

This version reports the three LongBench v1 English retrieval-QA subsets used in the main figure: **TriviaQA, HotpotQA**, and **2WikiMultihopQA**. These probe single-hop retrieval, multi-hop reasoning, and answer faithfulness under a common LongBench scoring protocol. The harness can also cap other text-QA benchmarks such as SealQA, FRAMES, and BioASQ for future sweeps, but they are not part of the main reported figure.

### Sampling

Each benchmark records the exact evaluated tasks. The three selected LongBench subsets use 200-question samples for ReAct and Plan-Execute and 120-question Plan-Evolve runs with prompt adjustment enabled. For cost-limited sweeps the harness caps a benchmark to the *first N questions in dataset order*—a deterministic, reproducible prefix, not a random independent sample. The selected tasks are persisted with the run metadata; comparisons are treated as paired only when the same examples are available for the strategies being compared.

### Scoring

Every answer is scored under multiple metrics so that verbose research-style outputs are compared fairly against short-answer leaderboards: a lenient *substring* match, *recall* of any gold alias, and three LongBench-style token-F1 variants over progressively tighter answer-span extractions (first sentence, first bold span, and the noun-phrase-tightened bold span used as the fair single-span estimate). We compare against published LongBench anchors at matched parameter scale where available.

### Future agentic-environment benchmarks

Benchmarks requiring live browser or desktop actuation—GAIA, WebArena / VisualWebArena / WebArena-Infinity, OSWorld, and AgentBench—stress brittle UI grounding, long-horizon state tracking, and execution recovery, but lie outside our current text-and-retrieval tool set (the agent has search and document-fetch tools, not a browser driver). We leave these, together with advanced-RAG retrieval benchmarks (BEIR, MIRACL), to future work.

## C Case Study Reviewer Evaluations

In the following, we reproduce the adversarial reviewer evaluations generated by ContinuumCellAgent for four Tier-1 biomedical case studies. Each evaluation covers the methodological strengths, integrity concerns, and biological validity of the agent-produced manuscript.

### C.1 PDAC (GPT-4 Plan-and-Execute)

#### Strengths

The agent completes a full analytical pipeline on real large-scale public data, downloading and processing GSE16515 (*N* =52) and GSE15471 (*N* =78) and producing verifiable differential expression outputs with an intact SHA-256 provenance chain. The reproducibility signal is statistically compelling: a cross-cohort directional concordance of 0.78 against a null expectation of 0.50, corroborated by 100% directional agreement with an external 300-gene reference set (*p* = 8 *×* 10^−179^), constitutes publishable replication evidence for a genuine research gap supported by prior literature. The agent also maintains exemplary transparency by proactively disclosing the substitution of a microarray workflow for the planned RNA-seq pipeline, clearly reporting the complete failure of enrichment analysis due to missing dependencies, and consistently distinguishing planned from executed steps.

#### Weaknesses

The phrase *robust replication* in the abstract overstates the strength of the findings; it should be revised to *directional agreement under a single analytical framework*. Biologically, the study yields no new discoveries: pathway enrichment failed entirely with no investigation into the cause, foreclosing mechanistic interpretation and leaving the core research objective unmet. The top-ranked differentially expressed genes are dominated by stromal markers, and stromal contamination is uncorrected, precluding any inference about tumor cell-intrinsic transcriptional changes. In sum, the manuscript is methodologically candid but scientifically incomplete, offering a demonstration of honest legacy microarray analysis rather than an independent scientific contribution.

### C.2 AD (Gemini Plan-and-Execute)

#### Strengths

The agent demonstrates a meaningful degree of methodological self-correction: Phase 2 appropriately rejects the causal claims regarding ARX made in Phase 1, as the AUPRC of 0.644 fell below the predefined threshold of 0.85, and a sensitivity analysis was conducted across 12 parameter combinations. The biological gap motivating this work is genuine: APOE4-associated microglial dysfunction as an early driver of AD represents an active area of inquiry, and integrating causal inference tools such as SMR and COLOC with single-cell data constitutes a real methodological gap. The Phase 2 manuscript is more transparent in its framing, explicitly labeling the model as a spatial autoregressive co-expression model rather than a causal inference system.

#### Weaknesses

The most critical issue is circular validation: the genes reported as causal drivers of AD were pre-specified in the data-generating process, so the analysis recovers its own input parameters rather than producing genuine biological discoveries. Additionally, the manuscript claims in Section 3.3 to have analyzed real ROSMAP and MSBB data, whereas the workspace contains only synthetic files of the form `rosmap_synthetic_*.csv` with no evidence of real data acquisition or processing. A second variant of the AD run further portrays a known method—incorporating aneuploidy score as a covariate in an NB-GLM design—as a novel framework, when this is standard practice in DESeq2 multivariate design and does not constitute a methodological innovation. As a consequence, the core biological conclusions are not credible under the available evidence.

### C.3 LUAD (Gemini Plan-and-Execute)

#### Strengths

The agent addresses the genuine and active research problem of aneuploidy confounding transcriptomic differential expression analysis. Employing a multi-factor DESeq2 design that includes CNV covariates is a known means of dealing with this issue, and the direction pursued by the agent is therefore reasonable in principle.

#### Weaknesses

The manuscript misleadingly presents synthetic simulated data as if they were genuine patient data. Additionally, a known method is described as novel: incorporating the aneuploidy score as a covariate in an NB-GLM is described as a novel framework, yet this is standard practice in the DESeq2 multivariate design and does not constitute a methodological innovation.

### C.4 Longevity (Gemini Plan-and-Execute)

#### Strengths

The agent maintains basic honesty in Phase 1 by explicitly stating in the manuscript that the data are 100% synthetic. The differential expression analysis based on *N* =30 synthetic samples using the Mann–Whitney *U* test with FDR correction follows standard methodological practice. The agent also correctly identifies genuine gaps in the longevity literature with largely reasonable descriptions.

#### Weaknesses

Several serious integrity issues arise in Phase 2. The GEO dataset accession numbers cited in the manuscript do not match those actually used in the code, a fundamental inconsistency that undermines reproducibility. Because the input data in Phase 1 are entirely synthetic, all 38 identified targets lack biological validity. The naked mole rat (NMR) dataset is further limited by an extremely small sample size of only 9 samples, which severely restricts any meaningful statistical inference.

